# Identification of Structures for Ion Channel Kinetic Models

**DOI:** 10.1101/2021.04.06.438566

**Authors:** Kathryn E. Mangold, Wei Wang, Eric K. Johnson, Druv Bhagavan, Jonathan D. Moreno, Jeanne M. Nerbonne, Jonathan R. Silva

**Author notes:** **Correspondence:**, Jonathan R. Silva, PhD, Washington University in St. Louis Campus Box 1097, 1 Brookings Dr. St. Louis, MO 63130.

## Abstract

Markov models of ion channel dynamics have evolved as experimental advances have improved our understanding of channel function. Past studies have examined various topologies for Markov models of channel dynamics. We present a systematic method for identification of all possible Markov model topologies using experimental data for two types of native voltage-gated ion channel currents: mouse atrial sodium and human left ventricular fast transient outward potassium currents. In addition to optional biophysically inspired restrictions on the number of connections from a state and elimination of long-range connections, this study further suggests successful models have more than minimum number of connections for set number of states. When working with topologies with more than the minimum number of connections, the topologies with three and four connections to the open state tend to serve well as Markov models of ion channel dynamics.

**Significance Statement:** Here, we present a computational routine to thoroughly search for Markov model topologies for simulating whole-cell currents given an experimental dataset. We test this method on two distinct types of voltage-gated ion channels that function in the generation of cardiac action potentials. Particularly successful models have more than one connection between an open state and the rest of the model, and large models may benefit from having even more connections between the open state and the rest of the other states.

## Introduction

Discrete state Markov, or state-dependent, models have been used extensively to probe the role of ion channel dynamics in generating the excitability of neurons(1), cardiac myocytes(2), and pancreatic beta cells(3,4). Markov models recapitulate channel dynamics by discretizing behavior into a series of states, with transitions between states governed by rate constants that often vary as a function of membrane potential(5). These Markov models are then inserted into cellular models to simulate action potential waveforms and frequency-dependent properties(3,6–8). For many types of ion channels, Markov model topologies describing their kinetic and voltage-dependent properties have evolved to reflect refined knowledge of channel behavior and functioning from decades of experimentation. For example, experiments have revealed multiple activation gates and inactivation states that span many time domains in some types of voltage-gated ion channels. States have been continuously added to existing Markov models at the single-channel and macroscopic current levels to improve their ability to account for this additional complexity (9–13). For the voltage-gated cardiac Na^+^ channel, for example, current Markov models reflect multiple stages of channel activation, deactivation, and inactivation from both closed and open states (14,15).

There is a rich history of progress in modeling macroscopic and single channel currents taking advantage of various topologies, or structures, of Markov models that reflect our understanding of channel dynamics (1,13,16–22). There have been also numerous studies on parameter identifiability and equivalence (18,23–28). In both cases, however, have explored a limited collection of topologies either for understanding the details of channel gating or recapitulating general channel dynamics. In 2009, Menon and colleagues surpassed previous efforts by exploring many model topologies through a genetic algorithm that theoretically optimizes model structure in addition to the rate parameters. Making random perturbations to optimize model structure, however, is challenging from an optimization point of view because the addition or removal of a state causes a large jump in the parameter landscape. By enumerating the unique channel Markov model topology search space, however, optimization may focus on rate parameterization of these unique structures that thoroughly cover the search space. Enumeration also allows for absolute ranking of structures in order of increasing complexity, so that through examining the performance of multiple topologies, one may estimate the complexity needed to recapitulate that specific dataset.

Systematically identifying various model structures is especially helpful given the range of goals in channel kinetic modelling(29). A model, for example, may need to reflect new structural information, new functional role(s) of channel interacting proteins (30–32) or, as in the CiPA initiative(33), massive amounts of electrophysiological data to simulate proarrhythmic effects of drugs. These types of studies may require a model that recapitulates molecular level detail precisely, i.e., for gating studies. Other types of studies, however, may only need a model that captures the principal dynamics of the channels, for example, in simulations of action potentials. By enumerating all possible model structures, we can suggest multiple structural candidates at different levels of complexity for validation of various types of data and complexities of datasets.

While human intuition laid a solid foundation for early ion channel models (34), there is a great need for a systematic, efficient method to identify possible Markov topologies given a specific experimental dataset. We present a systematic investigation into Markov model topologies examined incrementally in increasing complexity with two voltage-clamp datasets derived from analyses of cardiac fast transient outward (I_to,f_) potassium currents and rapidly activating and inactivating sodium (I_Na_) currents. Multiple topologies invite opportunities to understand how discretized states and rate constants come together to form a successful model of channel dynamics. This strategy also provides the opportunity to summarize topological features that work well for creating channel models.

## Results

Our initial aim was to count how many different topologies are possible for a Markov model with a given number of states. To accomplish this goal, we assumed a Markov model topology with one state designated as open for simulating current with the rest not strictly labeled as in Menon et al. (16). In terms of graph theory, this open state is called the root. By starting with the root (open state), we could then iteratively evaluate the connectivity of the other states (35,36). A challenge arose here, however, because topologies may appear to be unique by their numbering, even though the state labels are permutations. Thus, our counting algorithm needed to assess whether models were oriented uniquely, as opposed to simply being labeled differently. Using the 3-state topology space as an example, 36 permutations of single rooted topologies are possible (**Figure 1A**). **Figure 1B** then depicts the three topologies that are unique with respect to the root for clarity. A unique graph guarantees that the root, is oriented distinctly with respect to the other states. To generate this unique space for topologies with greater than three states, topologies of various sizes were tested for isomorphism (37) and only the unique topologies were retained (38). Parsing from single rooted topology permutations (36) to uniquely oriented state topologies (3) is depicted in **Figure 1C** along with the results of a similar analysis for topologies up to 10 states.

**Figure 1.**
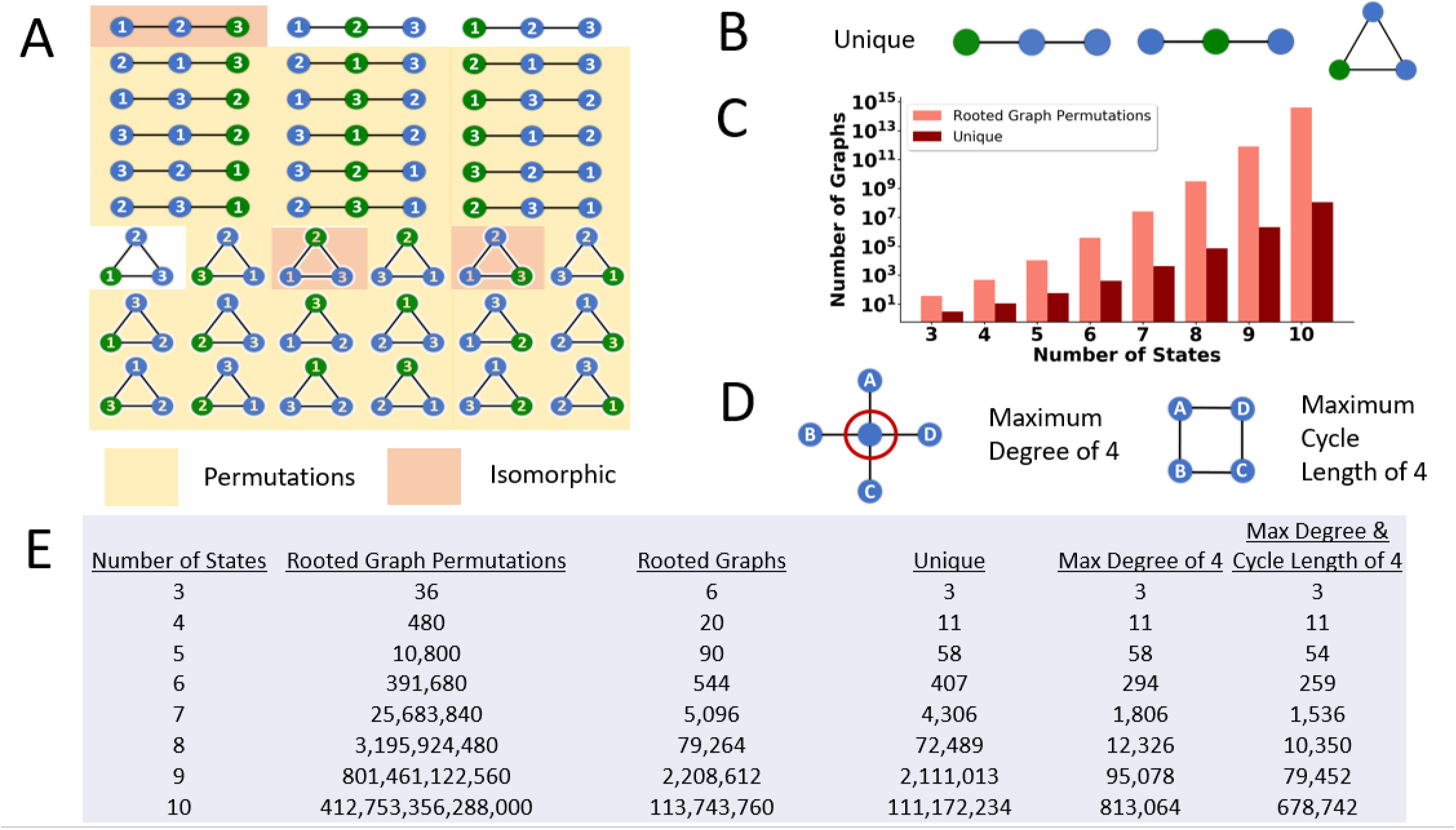
Reduction in model search space by enumerating unique topologies as a function of the states. **A)** All 36 rooted graph permutations of three states with highlighted permutations (yellow). Blue states are non-open while the open state(root) is colored green. Topologies highlighted orange and no highlighting represent the six possible rooted topologies with three states. Unhighlighted topologies are unique rooted topologies of size three. **(B)** while the orange shading represents the isomorphic topologies. A reduction from 36 rooted graph permutations to three unique topologies is depicted in **C**. **C)** Results of a similar graphical enumeration analysis for rooted topologies with 4+ states. **D)** Biophysically inspired restriction of the maximum degree in a graph to 4. Applying this restriction to the unique topologies results in a reduction in a model search space as enumerated in the table. After further restricting the maximum cycle length in a graph to size 4 after the degree restrictions, final graph counts are displayed in the table (**E**) as function of the number of states. **E)** Enumeration summary table of rooted graph permutations, rooted topologies, unique rooted topologies, and biophysical restrictions as a function of the number of states.

As can be seen in the table, this parsing dramatically reduced the model search space by orders of magnitude as the number of states increased, providing an upper bound on the topology search problem. However, the number of unique topologies was still quite large (on the order of millions for 9 and 10 states), and many topologies cannot plausibly be studied given current constraints on computational resources. Thus, we sought to reduce the number of topologies evaluated further by focusing on those that might be the most useful for modeling native ion channel behavior. We do note, however, that future efforts to explore the excluded models further might be feasible and appropriate as computational resources continue to increase.

To further reduce the number of topologies, the degree of a state, defined as the number of edges (connections) possible to other states given residency in a certain state, must be limited. We placed a moderate restriction of a maximum of a degree of four on a state (preventing one state from accessing many others) (**Figure S1A**). A state with a high degree implies that a given conformation of the channel has direct access to many different adjacent conformations, each with an associated rate. These rates would be increasingly difficult to identify experimentally as the number of connections increases.

Large cycles also introduce additional challenges as the topologies start to represent long-range connections. In other words, states that are far apart in the ion channel excitatory cycle such as “deep” (or especially stable) inactivated or closed states may be connected directly. Experiments suggest, however, that a sequence of distinct channel energetic conformations likely take place between these stable states (39,40). By retaining these long-range connections or cycles, the topology implies the distinct pathways may be bypassed. Like the maximum state connections, we set a moderate restriction of four on the maximum cycle length to create the focused model search space (**Figure S1B**). Together with elimination of isomorphic topologies, focusing the search on topologies that meet both biophysical restrictions **(Figure S1C)** yielded a reasonable number of models to evaluate given currently available computational resources, reducing the original space for a 10-state model from 10^14^ to a more reasonable 10^5^.

Our principal aim was to evaluate these unique topologies as Markov models of channel dynamics. The unique models with varying number of states (**Figure 1**) were sorted according to increasing numbers of free rate constants (proportional to the sum of the number of states and edges (connections) as a measure of model complexity (16). We utilized two canonical ion channel datasets to possible model structures needed to recapitulate channel dynamics: the rapidly activating and inactivating, voltage-gated cardiac sodium current (I_Na_) and the fast transient, voltage-gated outward potassium current (I_to,f_). In mammalian cardiac myocytes, I_Na_ is responsible for the upstroke of the action potential (41) while I_to,f_ contributes to early repolarization and, notably, is responsible for generating the early “notch” of the action potential that is prominent in epicardial left ventricular myocytes in many large mammals, including humans (42).

To evaluate the suitability of a particular topology to serve as a Markov model, we needed to find optimal rate parameters to capture the trends in the voltage-clamp data. Rate parameters were optimized using simulated annealing with adaptive temperature control (43) (equations 2–5). To minimize the dependence of optimal rate parameters of a unique graph on initial optimization conditions, multiple starts of simulated annealing were performed with initial rate parameters scaled according to a quasi-random Sobol sequence (44)(equation 9) to thoroughly explore the parameter space. We aimed to find rate parameters that captured the trends in the voltage-clamp data while also avoiding overfitting. Overfitting of a model means the model fails to represent the general trends in the data by focusing on fitting all experimental data perfectly (45). In other words, we wanted to maximize the chance the Markov models would successfully predict channel dynamics not necessarily included in our canonical datasets (maximize generalizability). Further, we also wanted to quantify how likely it was that overfitting was occurring during the optimization process and when, so that the process could be terminated. Our rate parameter optimization routine included a measure to halt optimization if likely overfitting occurs using methods borrowed from training neural networks (46). Experimental data were split into training and validation sets. The trajectory of the reduction in training cost was tracked (progress) periodically throughout the optimization along with the current validation set cost with respect to the minimum seen (generalization loss). **Figure S2A** shows representative training and validation cost trajectories during a model optimization. The training cost slowly, but steadily, decreases throughout the optimization while the validation cost varies erratically. If the validation cost first decreased but then consistently monotonically increased after some point in the optimization, there would be a clear stopping point to prevent overfitting. However, the spiking in the validation cost trajectory makes it difficult to identify a clear stopping point. To quantify where overfitting is likely occurring and, therefore, where to terminate the optimization, measurements of progress and generalization loss were computed at regular epochs **(Figure S2B)**. These measures give insight into how much better the model is recapitulating the training experiential data at the expense of recapitulating the validation set with the best fidelity. As described previously (46), there are many strategies using this ratio to decide to halt the optimization. For this study, we used the condition that three consecutive increases in the overfitting ratio results in early termination. The trajectories of the progress and generalization loss measures demonstrate that this ratio allows enough flexibility for the validation cost to fluctuate throughout the optimization until a validation cost minimum is reached with termination shortly thereafter.

We also considered minimizing model solution stiffness while optimizing rate parameters for a given unique topology. Systems of differential equations are considered “stiff” when the derivatives of the function are large near the solution, requiring very small time steps to be taken for solution stability (47). Implicit differential equation solvers can be utilized to solve stiff systems more efficiently, although these are more computationally demanding than explicit solvers and often require specification of the Jacobian. The long-term goal here is to be able to incorporate the optimized channel Markov models into cellular and tissue models of membrane excitability. When scaling up to the cell or tissue level, a given ion channel model may be solved thousands and thousands of times. Thus, it is crucial that computational solving time is considered when creating the individual ion channel models. To quantify the stiffness of each model solution, the condition numbers of the transition matrices were estimated at various membrane voltages (48,49) (equation 7). Estimated reciprocal condition numbers larger than a threshold were averaged and proportionally contributed to the model cost as the stiffness penalty (equation 8).

We first present data from multiple optimizations runs with the overfitting and stiffness penalties for the human ventricular I_to,f_ dataset. **Figure 2A** displays the frequencies of optimization iterations completed as a function of increasing free rate constants while optimizing with overfitting and stiffness penalties. Iterations completed from multiple optimizations with differing starting conditions of each model are displayed. The relative weights of the black dots represent the frequency of maximum optimizations completed. As the number of free rate constants increases, most optimization runs reach the maximum number of allowed iterations (large black clusters). However, the intensity of the trailing black dots, which represent the number of optimization iterations completed before being terminated early, increases as well. This result supports the notion that overfitting becomes more problematic as the model complexity increases (24). **Figure 2B** depicts the associated normalized costs for the model populations after the optimization iterations specified in **Figure 2A.** The great spread in normalized costs illustrates the importance of running optimizations multiple times when estimating the absolute minimum cost of models with varying complexities. Of note, extremely high normalized costs result from especially poor optimization starting conditions.

**Figure 2.**
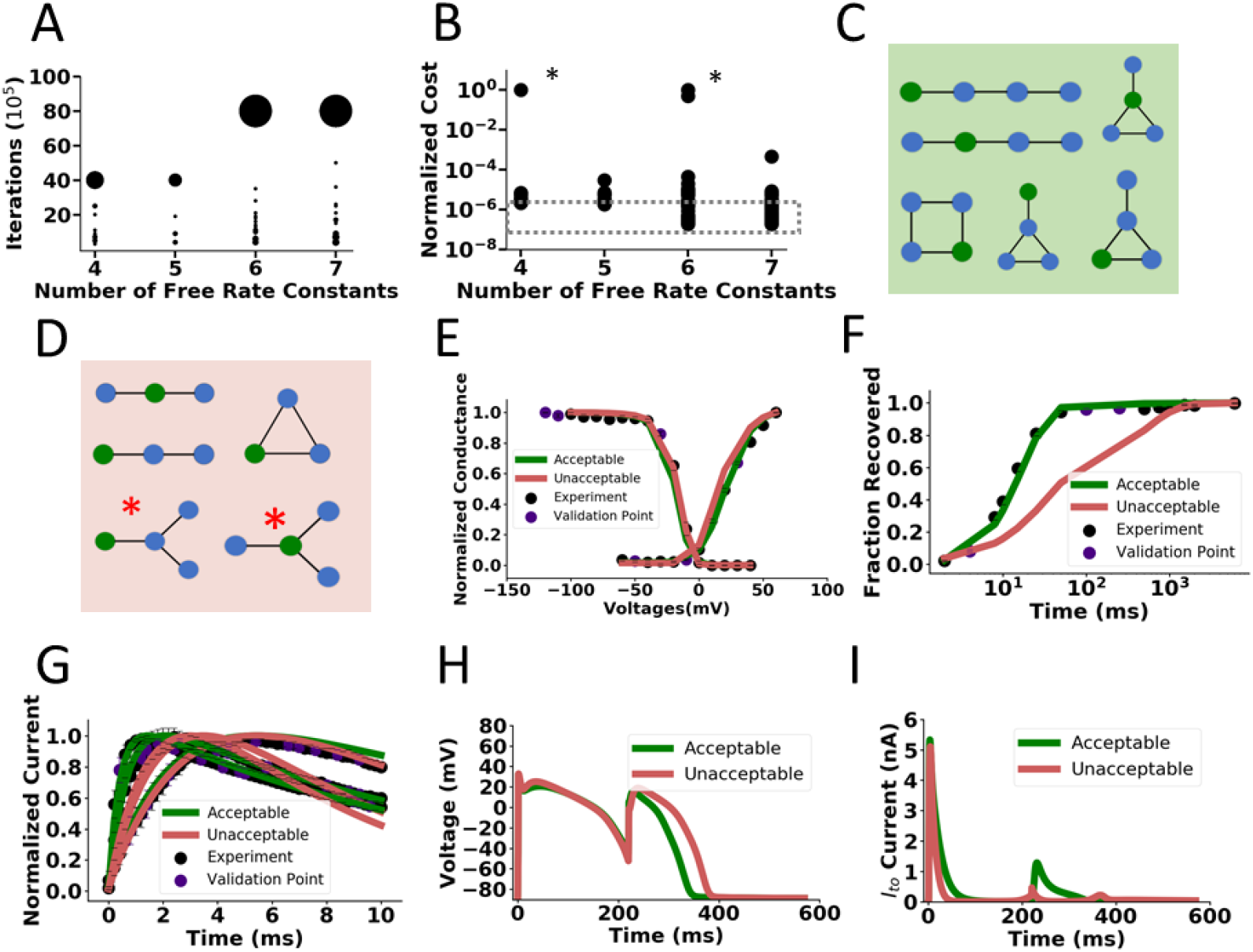
Identification of possible structures for the I_to_ dataset. **A)** Optimization iterations completed as a function of free rate constants. Weighting of dots represent the frequency of models completing a specified anumner of iterations. Most models complete the generous maximum iteration limit (40,000,000) with 4 and 5 free rate parameters while few models complete less iterations due to early stopping from the overfitting criterion. Models with 6 and 7 free rate parameters may run for longer (maximum 80,000,000 iterations displayed) while a greater fraction of models are terminated early due to overfitting criterion. Data points include the multiple starting conditions for each model to reduce the dependence of the minimum solution on initial conditions **B)** Distribution of normalized costs for each model with multiple starts after completing optimization iterations as depicted in **A**. Especially bad Sobol starting conditions are asterisked. The absolute minimum costs are outlined in the dashed grey box. The point of diminishing returns is at 6 and 7 free rate constants. **C)** Topologies producing acceptable fits and **D)** unacceptable fits for I_to_. Acceptable fits include models with four states and sufficient connections to separate the activation and inactivation domains. **E)-G)** Representative models fits for steady state activation, inactivation, recovery from inactivation and current traces for models in the acceptable and unacceptable model categories. Unacceptable models generally have very slow recovery from inactivation. **H)** Simulated action potentials with representative acceptable and unacceptable I_to_ currents under a S1-S2 protocol. **I)** Corresponding simulated I_to_ currents under the S1-S2 protocol in **A**. Acceptable currents (green) and unacceptable currents (red) most differ in magnitude at around 200 ms into the S1 action potential with much less unacceptable I_to_ current. This corresponds with the slow recovery from inactivation depicted in **F)** where at ~200 ms the acceptable models have fully recovered while unacceptable models are only half recovered.

Focusing just on the absolute minimum costs seen across the model populations reveals that the minimum cost trends downward as complexity increases: models with six and seven free rate constants produce minimum costs on the order of 10 times less than models with four free rate constants. Tracking this absolute minimum cost as the number of free rate constants increases reveals a point of diminishing returns. The absolute minimum cost decreases appreciably when comparing model populations with four, five, and six free rate constants. However, there is hardly any change when comparing models with six and seven free rate constants. **Figure S3** displays the various stiffness penalties as a function of model complexity. The smallest stiffness penalty seen across all optimization starts levels out for model topologies with six and seven free rate constants, as does normalized cost. This point of diminishing returns in absolute minimum cost and stiffness penalties at six free rate constants suggests that this complexity may be optimal to model the canonical dynamics of I_to,f_ in this specific dataset. Models with six to seven free rate constants have the potential to prioritize good fidelity fits to voltage protocols while minimizing overfitting potential. **Figure S4** displays the topology of an example model for the I_to,f_ dataset with six free rate constants along the fits to the voltage protocols.

Our next aim was to categorize all I_to,f_ model topologies studied as acceptable or unacceptable, based on the minimum cost seen across all optimization starts. A model with a minimum cost no greater than 300% of the absolute minimum was deemed “acceptable.” These topologies are displayed in **Figure 2C** and highlighted in green. Models that consistently produced poor voltage protocol fits are displayed in **Figure 2D** and highlighted in red. These unacceptable models have four to five free rate constants and consistently produce higher normalized cost values, as illustrated in **Figure 2B**. Representative model fits from the acceptable and unacceptable model categories to the voltage-clamp protocols are displayed in **Figures 2E-G** and colored accordingly. Unacceptable models tended to produce fits with slow recovery from inactivation **(Figure 2F)** with little impact on the other protocols **(Figures 2E,G)**. **Figure S5** tracks state occupancy as function of time during the recovery from inactivation protocol. Acceptable models have enough complexity to recapitulate the slow timescale at steady state and the faster timescale during recovery at −70 mV. Unacceptable models show slow recovery from inactivation because the rates cannot be sufficiently fast during recovery while also fitting steady state conditions.

We also included the modeled I_to,f_ into a human ventricular myocyte action potential model (50) under a S1-S2 pulse protocol that simulates repetitive excitation (see Methods) to validate our categorization of acceptable and unacceptable models based on cost. An S2 stimulus given at around 200 ms into the S1 action potential revealed that unacceptable models can lead to a longer action potential duration **(Figure 2H)**. Analyzing the corresponding currents revealed that, at 200 ms into the S1 action potential, the magnitude of I_to,f_ generated was much lower in the unacceptable model compared to the representative acceptable modeled I_to,f_. **(Figure 2I)**. This result is in accordance with the lagging fraction of recovered channels for the unacceptable models at 200 ms as depicted in **Figure 2F**. As the magnitude of I_to,f_ influences the notch and plateau potentials, which will secondarily impact calcium entry and excitation-contraction coupling (51), the deficiencies in the simple models could result in inaccurate cellular and tissue level predictions. Analyzing the modeled currents under this S1-S2 protocol to stimulate the impact of changing heart rate reveals the precise window of time over which the inactivation and incomplete recover from inactivation of I_to,f_ channels could manifest itself at the cellular level. The overly simplistic models, with complexities below 6-7 free rate constants, fail to capture the full dynamics of I_to,f_ when simulating rate-dependent effects on action potential waveforms.

We then repeated this analysis on an available (mouse atrial myocyte) I_Na_ dataset to find possible models for a channel with more complex dynamics so that we could explore more complex topologies. The cardiac sodium current is more complex than I_to,f_ because of the fact that channel activation and inactivation are both very fast (41). As before with I_to,f_’ we aimed to sort the studied model topologies in order of increasing complexity into the acceptable and unacceptable model categories, as defined before, based on cost (**Figures S6-S7**). While examining the model fits to the individual voltage protocols helped classify acceptable and unacceptable topologies for the I_to_ dataset, acceptable and unacceptable I_Na_ models did not display a severe protocol fitting deficiency, meaning one of the protocol fits was consistently poor (**Figures S7E-G**). Unsurprisingly, when validating the simulated acceptable and unacceptable I_Na_ currents in the action potential (52), there were no appreciable differences in morphology (**Figure S7H**).

Because the I_Na_ acceptable and unacceptable models could not be distinguished by the representative individual protocol fits, we then repeated our I_Na_ analysis on a previously published I_Na_ dataset, generated in HEK-293 cells, that includes slow and intermediate components of recovery from inactivation (53–55). We predicted that a more complicated recovery from inactivation protocol would require greater model complexity to fit all voltage protocols, so that topologies with few free rate constants would fail to reproduce all kinetics. **Figures 3A and 3B** show representative acceptable and unacceptable model topologies after rate parameter optimization confirming this hypothesis. The models needed at least eight free rate constants to fit this complex dataset **(Figure 3A),** while sparsely connected topologies and those with fewer than eight free rate constants **(Figure 3B)** failed to reproduce all protocols with good fidelity. Representative unacceptable and acceptable model fits to individual voltage protocols are shown in **Figures 3C-F**. **Figure 3E** depicts the more complex protocol of recovery from use dependent block (RUDB). The repetitive voltage steps generating RUDB (see Methods for protocol) allows for the slower and intermediate components of recovery from inactivation of an I_Na_ model to be parameterized. Unacceptable models show slow recovery from fast inactivation **(Figure 3D)**, poor model fidelity to the intermediate and slow timescales of recovery from inactivation under RUDB **(Figure 3E)**, and poor voltage-dependent inactivation kinetics **(Figure 3F)**. This result means that it is quite difficult to parameterize simpler models to capture accurately various types of recovery from inactivation along with other voltage protocols. We then repeated the action potential validation analysis on the HEK I_Na_ dataset categorization of acceptable and unacceptable models based on cost. Incorporation of modeled currents into the action potential (52) shows that unacceptable I_Na_ HEK models tend to produce action potentials that fail to repolarize, while representative acceptable models successfully repolarize **(Figure 3G).** Plotting the simulated I_Na_ reveals that representative unacceptable models have abnormal gating into the action potential (late component) that hinders action potential repolarization **(Figure 3H)**.

**Figure 3.**
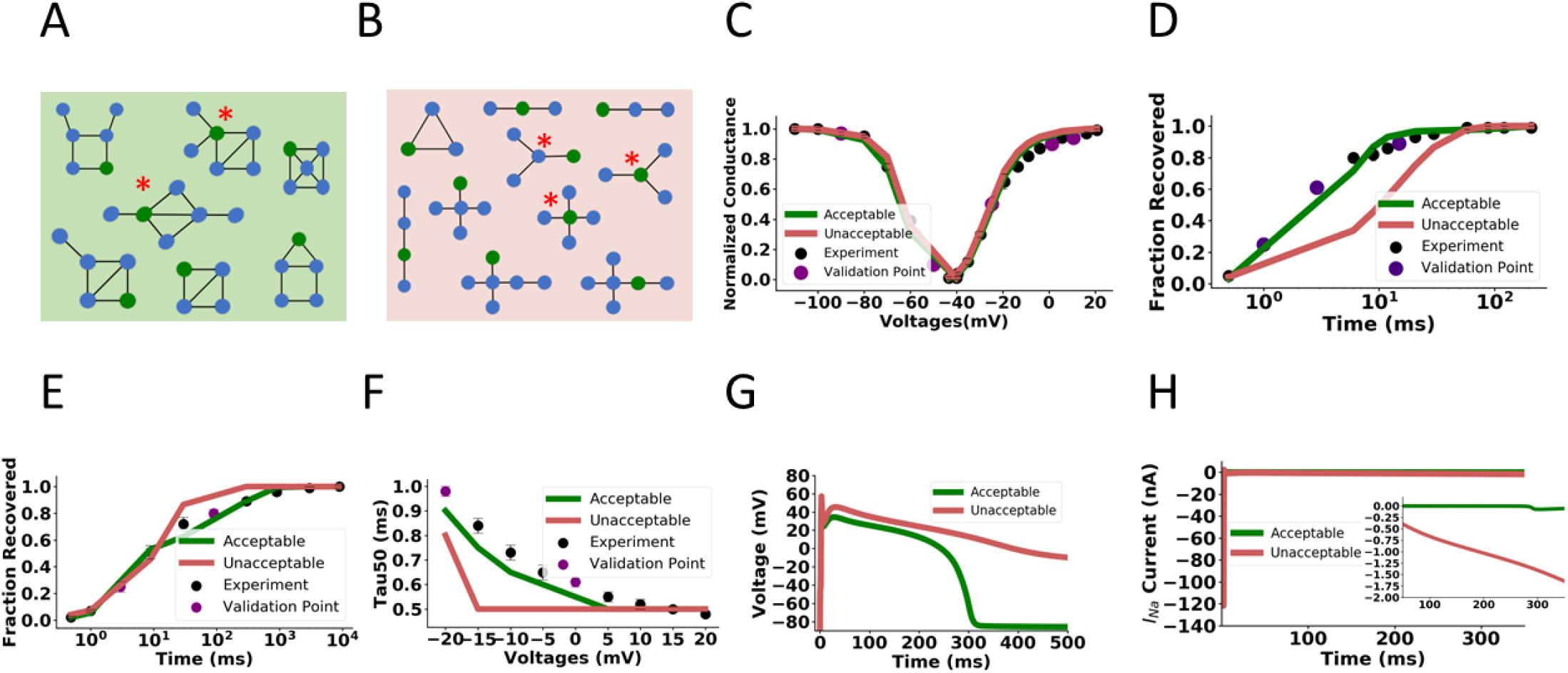
Identification of possible model structures for the I_Na_ HEK dataset. **A)** Representative acceptable model topologies of the more complex HEK I_Na_ dataset. Topologies have at least 8 free rate constants. **B)** Representative unacceptable model topologies. Topologies have fewer than 8 free rate constants or are sparsely connected. **C)** Representative acceptable and unacceptable model fits to steady state activation and inactivation **D)** Representative fits to fast recovery from inactivation **E)** Representative fits to recovery from use dependent block which includes timescales of fast, intermediate, and slow recovery from inactivation. **F)** Representative model fits to the time constant of 50% inactivation of the peak sodium current **G)** Simulated action potentials with representative acceptable and unacceptable I_Na_ modeled currents. Unacceptable models have varying degrees of late I_Na_ current) **H)** Corresponding representative acceptable and unacceptable I_Na_ currents during the action potentials in **G** with a magnified inset highlighting the late I_Na_ current.

Up until this point, we reported the minimum cost seen across all optimization starts when describing a model as acceptable or unacceptable. In Figure 4, we report the end performance of each optimization start (20 displayed) for each topology in addition to all low-cost model solutions in the reported optimization history. This presentation reflects the diversity in rate parameter parameterizations (i.e. model solutions) during and after multiple optimizations. For example, for a given topology there may be vastly different sets of parameters that produce a model with an acceptable cost. **Figure 4A** summarizes the performance of all studied I_to,f_ model topologies to serve as Markov models when sorted by number of states and connections. Acceptable and unacceptable labels correspond to the action potential validation performance as depicted in **Figure 2H**. However, 23 out of 167 low cost model solutions on topologies depicted in **Figure 2C,** failed to recapitulate the increased I_to,f_ evident using the S1-S2 protocol. We labeled those topologies as “tentative” to reflect the fact that, despite good fits to the voltage-clamp protocols, these two models failed to perform like the other acceptable models in the action potential simulations. In **Figure 4A**, four-state models with four edges were the most likely to produce acceptable models, but notably a linear four-state model with three edges was the minimum cost model for the I_to,f_ dataset **(Figure S4)**. **Figure 4B** displays that topologies with lower root degree (fewer open state connections) tend to be more successful as Markov models. This observation makes sense given that topologies with lower root degrees with few states result in a “spoke” layout (see asterisked models in **Figures 2D and 3B**). These “spoke” topologies with a central open state require careful rate parameterization to prevent “bursting” of the channel over time at various membrane potentials. This translates into a more difficult optimization problem, and so our optimizer struggles to find a satisfactory solution given our finite optimization limits.

**Figure 4.**
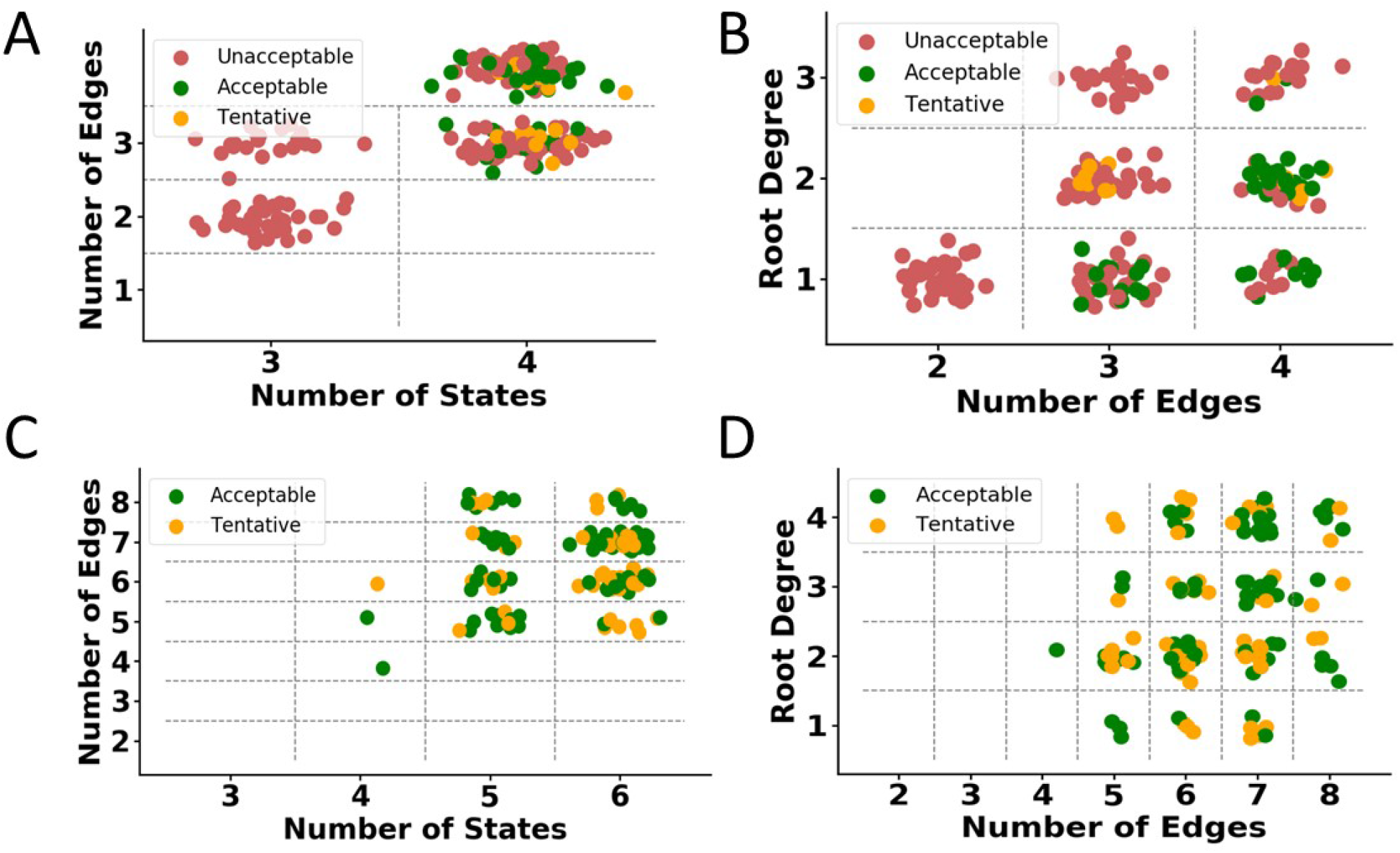
Summary of Markov model performance for all topologies studied **A)** and. **B)** Performance summary of all I_to_ topologies studied. Unacceptable and acceptable labels are as previously defined. Tentative yellow topologies produced acceptable voltage protocol fits but did not perform as other acceptable models in the action potential validation. In **A)** more states help topologies find tentative and acceptable solutions. **B)** Topologies with lower root degrees (open state connections) tend to create acceptable models **C** and **D)** Performance summary of all I_Na_ topologies studied. Unacceptable and acceptable labels are as previously defined. Tentative yellow topologies produced acceptable voltage protocol fits but did not perform like other acceptable models in the action potential validation (different degrees of repolarization failure). **C)** Topologies with more than the minimum number of edges tend to yield more tentative or acceptable Markov models. **D)** As edges increase, topologies most benefit from higher root degrees (more open state connections) to serve as successful Markov models. Panels C and D do not show unacceptable models for clarity.

By analogy, **Figures 4C and 4D** summarize the performance of all model topologies studied across all optimization starts and history when training on the HEK I_Na_ dataset. For clarity, only the acceptable and tentative models are displayed in **Figures 4C and 4D** while **Figure S9** includes unacceptable models as well. As before, acceptable, and tentative labels correspond to the performance of each topology during action potential validation in **Figure 3H**. Out of the total of 169 acceptable model solutions based on cost, 68 are labeled “tentative” to reflect their differing behaviors, compared with the other acceptable models, in the action potential simulations. When sorting based on number of states and edges, topologies with more than the minimum number of edges tend to serve as successful Markov models **(Figure 4C)**. In other words, sparsely connected topologies do not have the complexity to recapitulate channel dynamics. When sorting based on total topology connections versus root degree in **Figure 4D**, as the number of edges in a topology increases, higher root degrees aides in creating a successful Markov model. Once a topology has seven connections, for example, the topology likely has six states, so open state connections may still be incorporated in a cycle (sample topologies asterisked in **Figure 3A).** This arrangement allows the topology to recapitulate more complex dynamics. Thus, many open state connections are not automatically detrimental for topologies with more complexity.

Taken together, sparsely connected topologies tend not to serve as successful Markov models. When topologies have five states or less, more open state connections result in harder to parameterize “spoke” topologies. However, when topologies become complex enough (many connections or states), this is not an inherent detriment to the topology. A root then may have many connections to the rest of the model and still be incorporated in a cycle, for example, so the topology has inherently more capacity to represent more complexity. This thinking suggests potential filters to further parse the model space beyond degree and cycle length restrictions. For topologies with three, four, or five, states (2–3) open state connections should be emphasized while larger models will benefit from a variety of open state connections. Given limited computational resources, topologies with more than the minimum number of connections should be prioritized. For example, in the seven-state model space, there are 1483 topologies after applying restrictions on state connections and cycle length. (**Figure 1E**). If topologies with minimal connections (6–7) were excluded, another 166 topologies could be further parsed.

## Discussion and Future Directions

We present a robust, systematic method to identify all possible model topologies for simulating current, given an experimental dataset of canonical ion channel dynamics. The routine moves through various topologies in a stepwise fashion as a function of the number of free rate constants. By examining the diminishing returns in model cost, one may visualize the complexity of various topologies to best balance between model solution fidelity, overfitting (24), and stiffness for the specific experimental dataset. Depending on the goals of the kinetic modeling study undertaken, the user may still wish to utilize a topology with more or less complexity identified in this routine. In these cases, we provide an organized framework for users to validate topologies for their specific goals. We demonstrate the robustness of this methodology by modeling two voltage-gated currents: fast transient outward (I_to,f_) potassium currents and rapidly activating and inactivating sodium (I_Na_) currents.

Using the formulation of the single tracked open state as in Menon et al. (16), we were able to enumerate the model topology search space. This exhaustive enumeration ensures complete coverage of the model search space, rather than relying on random perturbations or limited collections of various model topologies during optimization. As illustrated in **Figure 1**, this enumeration is crucial when working with topologies with six states and greater because of permutations. Enumeration also allows for topologies to be evaluated systematically in terms of model complexity, which is critical for identifying the amount of complexity available in possible structures for an experimental dataset. Through restrictions on state connections and elimination of long-range connections, we were able to further parse this model search space given decades of biophysical experimental insight. This study further suggests an additional filter that eliminates topologies that are sparsely connected. It may also prove fruitful to emphasize topologies with lower root degrees (fewer open state connections) when working a few states but higher root degrees (many open state connections) (up to four) in topologies with seven states and higher.

Optimizing models with greater number of parameters does not easily lead to a “perfect” model with a normalized cost of zero. With more computationally expensive starts and larger maximum simulation iterations without overfitting prevention, we may expect to see the cost function decrease to zero given infinite optimization time. However, given constraints on time and computing resources, we present the best costs at least 100,000 iterations beyond a change of 20% or less in cost (unless terminated early for overfitting). We see a point of diminishing returns in the normalized cost function and suggest optimal complexity, based on the dataset.

Grouping acceptable and unacceptable models of each dataset allows us to begin to answer the question of what makes a suitable topology for Markov models of ion channel dynamics. The I_to,f_ dataset showed that three-state models are not complex enough to recapitulate dynamics, but four-state topologies without cycles can successfully do so. The example minimal four-state linear model for I_to_ illustrated in **Figure S4A** and **S5C** does not contain a cycle between open, hypothetically closed and inactivated states commonly used to model voltage-gated ion channels (14,15). Ion channel Markov models need not always conform to human intuition of the underlying structural mechanisms to reproduce a dataset with good fidelity. A modeler has great power in determining how much complexity is needed for the computational problem at hand (56). More complex models, for example, may be appropriate in studies focused on exploring channel gating precisely. In studies that require attention to fast computations at the tissue level, a simpler, less stiff model, may be the most appropriate.

The example models of the simpler I_Na_ and the more complex I_Na_ datasets are more consistent with mechanistic intuition derived from consideration of experimental data. The example topology for I_Na_ in **Figure S8A** contains a cycle, which is consistent with prior cyclic models of ion channel excitability with connections between activated, inactivated, and closed states. Acceptable models depicted in **Figure S6C and Figure 3A** contain at least one cycle. Unacceptable I_Na_ models tend to be sparsely connected, thus not able to include cycles (aside from the three-state cyclic model), which leads to insufficient complexity to model fast channel dynamics. Performing analyses of the state probabilities during voltage protocols after using our procedure allows one to gain insight into the mechanisms of the model and discover surprising topologies that successfully recapitulate the protocols.

Validating the cost thresholds of acceptable and unacceptable current models when these are incorporated (with multiple other ionic conductances) into action potential models allows one to begin connecting how fits to voltage-clamp data will result in electrophysiological differences at the cellular level. In most cases, the action potential morphology may be predicted based on the cost of the model. In the case of I_to,f_, the kinetics of recovery of the channels from inactivation alters the early and late phases of the action potential consistent with marked frequency-dependent effects. The poor protocol fits in the HEK I_Na_ dataset, however, commonly resulted in severe action potential repolarization abnormalities. This result might have been expected, given that I_Na_ is solely responsible for the upstroke of the action potential in atrial and ventricular cells, while I_to_ is one of many currents responsible for repolarization.

However, we also found that 23 low cost I_to,f_ models solutions (out of 167 acceptable model solutions based on cost) and 68 HEK I_Na_ models (out of 169 acceptable model solutions based on cost), did not perform as well as the other acceptable, low cost models when incorporated into the action potential model. This discrepancy is not unknown in the ion channel modeling field. Ion channel modelers have traditionally specified one model topology to represent a complex dataset. If the optimized rate parameters for the topology failed to produce an action potential, the modeler would blame the topology and adjust the states and connections accordingly. This study reports on the performance of all possible model topologies, however, so this discrepancy becomes more apparent. Because optimized rate parameters can lead to satisfactory training data fits, yet behave differently in the action potential, it highlights whether a training dataset contains enough information to reliably constrain all model rate parameters consistently. This problem of parameter identifiability has been thoroughly discussed and quantified in ion channel and cellular models of excitability (18,23–28). Validating model performance in action potential simulations serves as an additional test to detect early signs of parameter unidentifiability and overfitting. Repeating the analysis with different protocols may further reveal which parameters are most critical for successful action potential generation, depending on the simulation to be performed and those most likely to suffer from parameter unidentifiability. These insights may suggest refined voltage protocols for these channels as previously done with hERG (57) to more efficiently train model parameters with more certainty.

These acceptable model solution totals for both datasets included model solutions based on the costs found at the end of each optimization and at intermediate recorded timepoints. For I_to,f_, there were 144 acceptable model solutions for the 11 distinct topologies studied. In the HEK I_Na_ dataset, 167 distinct topologies were studied, but only 169 acceptable model solutions were found across all optimizations. Thus, our optimization method found a relative abundance of acceptable solutions for the I_to,f_ dataset compared to the HEK I_Na_ dataset. This difference is likely attributable to the more complex nature of the HEK I_Na_ dataset with the faster kinetics the RUDB protocol, for example, which would require precise parameterization. When tracking the cost over time in the optimization for both datasets, the I_to,f_ optimization proceeded smoothly with a variety of acceptable solutions while the HEK I_Na_ optimization trajectory was substantially more jagged.

There are limitations to the approach presented, and these may be addressed by future work. We randomly set aside 20% of experimental data for validation. We certainly anticipate more sophisticated validation sets will be used in future studies. Additional validation data will prove useful, along with an analysis of the best data to serve as validation, in future iterations of this systematic method to minimize overfitting in models of higher complexity beyond those studied here. As depicted in **Figures 2A** and **S6A**, tracking three consecutive increases in the ratio of the generalization loss and progress successfully truncated optimizations, but future studies that explore other validation quantification cutoffs as the optimization problem evolves (24) would be valuable. We use the nonlinear optimization technique of simulated annealing with adaptive temperature control in this study, but more recent algorithms, such as particle swarms (17), may prove to be faster or more accurate. We anticipate other additions to the simulated annealing core routine could be helpful, such as adaptive simulated annealing (58) or differential evolution (59) in future studies.

This work provides a framework to identify multiple topology models for canonical ion channel kinetics. By providing open-source code of the computational routine, others may apply this routine to their biophysical systems. These populations of possible model topologies may suggest further experiments for validation of their behavior and may even elucidate more refined voltage-protocols for training the models. They will also allow for connections of the mechanistic underpinnings of the channel through analysis of their topology, rate constants, and state probabilities during voltage protocols. This intuition will prove invaluable when building models based on these topologies in future studies that recapitulate structural and drug interaction data.

## Methods

### Generation of nonisomorphic rooted unlabeled, connected topologies

Nonisomorphic (unique) topologies were generated using nauty (http://users.cecs.anu.edu.au/~bdm/nauty/) a C-based graph isomorphism testing routine. All connected, unlabeled topologies were generated with the specified number of states and with up to the maximum number of edges. (N(N-1))/2. The states in the topologies were then colored in two shades (root and nonrooted) and then imported into a routine to test for isomorphism through canonical labeling(37). These topologies were then imported and parsed in Python using the NetworkX package based on maximum degree and cycle length. After recording the degrees of all nodes in the graph, topologies were retained if all nodes did not exceed the maximum node degree restriction. Maximum cycle length was used to parse based on long-range connections in graph (see Results for an explanation for long-range connections). The routine ‘cycle_basis’ was used to identify the cycles in the graph and only topologies with cycles not exceeding the limit were retained.

### Electrophysiological Recordings

Voltage-clamp recordings for the simpler I_Na_ dataset were obtained from mouse left atrial myocytes at room (20-22°C) temperature. Experiments were performed using an Axopatch 1D (Molecular Devices) or a Dagan 3900A (Dagan Corp) patch clamp amplifier interfaced to a Dell microcomputer with a Digidata1332 analog/digital interface and the pCLAMP10 software package (Molecular Devices). For recording whole-cell Na^+^ currents, pipettes contained (in mM): 5 NaCl, 90 CsCH_3_O_3_S, 20 CsCl, 1 CaCl_2_, 10 EGTA, 10 HEPES, 4 MgATP, 0.4 Tris-GTP at pH 7.2, 300-310 mOsm. The bath solution contained (in mM): 20 NaCl, 110 TEACl, 10 CsCl, 1 MgCl_2_, 1 CaCl_2_, 10 HEPES, and 10 glucose, pH 7.4, 300-310 mOsm. See reference (60) for the experimental methods used to acquire the I_to_ dataset and references (53,54) for those used to acquire the I_Na_ HEK dataset.

### Evaluation of Unbiased Topologies to Recapitulate Canonical Ion Channel Dynamics

A biophysically focused model topology was trained on I_to_ human left ventricular myocytes, I_Na_ voltage-clamp protocols in atrial mouse myocytes and I_Na_ voltage-clamp protocols in HEK cells. Model rate constants were guaranteed to satisfy microscopic reversibility as outlined in Menon et al.(16) with a vector-valued voltage function as in Teed et al.(20):

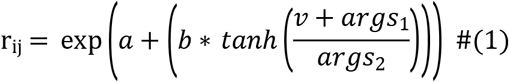

where *r_ij_* is the rate from state *j* to state *i*, *v* is voltage in (mV), *args*_1_ and *args*_2_ are optimized parameters as part of the vector valued voltage function (20), Values of *a* and *b* for each rate constant are listed in the SI for I_to_ and I_Na_ HEK dataset example models. As mentioned in Menon et al.(16), the rate constants are exponential functions of steady state occupancies and rates (one-ion symmetrical barrier pore model) (61).

Parameters are optimized using an improved simulated annealing routine. The improved simulated annealing routine included multiple noninteracting chains to effectively “parallelize” and thereby speed the optimization process (62) and an adaptive temperature control scheme (43). This temperature scheme begins at the lowest threshold temperature and is slowly incremented proportionally to the number of worse solutions encountered. When a “new” best solution is found, the temperature returns to lowest threshold. This scheme prevents the optimization from getting “stuck” in local optima:

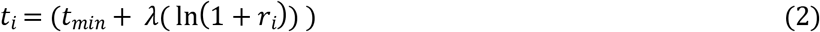

where *t_i_* is the current temperature is iteration *i*, *t_min_* is the minimum starting temperature, λ is the temperature control parameter and *r_i_* is determined by the change in the best cost, ΔC, at iteration *i*:

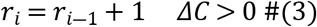

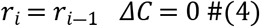

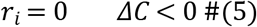

The cost function of the optimization was proportional to the sum of squared differences between each experiment and data value normalized by the experimental value plus a stiffness penalty:

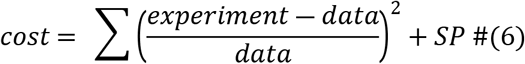

A model stiffness penalty proportional to each optimized model’s reciprocal condition number (1-norm) of the transition matrix, *A*, at varying voltages {1…N} was added to each model’s cost:

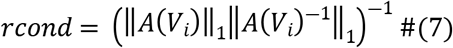

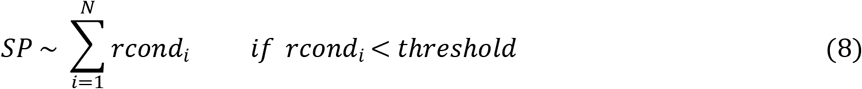

This penalty preferentially selects models that do not require extremely small, computationally expensive, time steps when incorporated into cellular and tissue level excitability simulations. To lessen the risk of the optimal rate parameters depending on the initial starting conditions, the optimization included multiple starts (at least 20) with a quasi-random (Sobol) representation of the parameter space (44). Quasi-random sequences increase coverage of the parameter space in each dimension (i.e. each free parameter is a dimension). Each sequential quasi-random sequence of dimension *D* is mapped to the specified ranges for the *i^th^* parameter value:

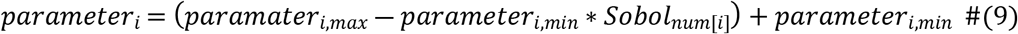

Each start ran for at least 100,000 iterations beyond no change in 20% of cost unless terminated early for overfitting prevention. Because we expected time to convergence would depend on model complexity, the maximum optimization iterations allowed were periodically increased for convergence. To prevent overfitting in the simulated annealing optimizations, the cost function only applied to model values outside each of each data point’s SEM. Quantitative metrics were also introduced to determine when to appropriately halt the optimization (46). The measure of generalized loss was calculated periodically throughout the optimization to quantify how the current validation error (given the current best solution) compares to best seen so far. Another measure of progress quantifies how fast the cost function has been decreasing the last k iterations. Three sequential increases in ratio of the generalization loss progress results in early termination to prevent overfitting.

#### Training set of voltage-clamp protocols

##### Steady state activation

I_Na_ HEK dataset: Steady-state probabilities were found at −100 mV. For voltages ranging between −45 mV to 20 mV, peak current was recorded after a step depolarization for 25 and normalized to maximum. I_to_: Steady-state probabilities were found at −70 mV. For voltages between −60 and 60 mV in increments of 10 mV, the peak current was recorded after a step depolarization for 50 ms and normalized to the maximum.

##### Steady state inactivation

I_Na_ HEK dataset: Steady-state probabilities were found at −100 mV. A conditioning pulse at voltages between −110 mV to 40 mV in 10 mV increments was applied for 500 ms. The peak current was then recorded after a test pulse at −10 mV for 25 ms and normalized to the maximum. I_to_: Steady-state probabilities were found at −70 mV. Each preliminary voltage step in increments of 10 mV between −120 and 40 mV was held for 200 ms. The peak current was then recorded after a test pulse at 40 mV for 50 ms and normalized to the maximum.

##### Recovery from inactivation

I_Na_ HEK: Steady-state probabilities were found at −100 mV. A depolarizing pulse at −10 mV for 500 ms was applied, followed by a hyperpolarizing pulse at −100 mV ranging between 0.5-210 ms. Peak current current was then recorded and normalized after a pulse at −10 mV for 25 ms. I_to_: Steady-state probabilities were found at −70 mV. A depolarizing pulse at 40 mV for 500 ms was applied, followed by a hyperpolarizing pulse of −70 mV of variable time intervals (2-6000 ms). Peak current was then recorded and normalized after a pulse at 40 mV for 100 ms.

###### Recovery from Use Dependent Block (I_Na_ HEK only)

Steady-state probabilities were found at −100 mV. A pulse train of a depolarization at −10 mV for 25 ms at 25 Hz was repeated for 100 pulses. A hyperpolarizing pulse at −100 mV for variable recovery intervals was applied for between 0.5-9000 ms. A test pulse followed at −10 mV for 25 ms and peak current was normalized to the maximum.

##### Normalized Current Traces

(1_to_ only): Steady-state probabilities were found at −70 mV. Following a step depolarization to 20 and 60 mV for 10 ms, the normalized current was recorded at intervals of 0.2 ms.

##### Deactivation Time Constants (I_Na_ only)

Steady-state probabilities were found at −120 mV. Following recording the peak current after a depolarizing pulse at −20 mV for 5.0 ms, a hyperpolarizing voltage between −110 mV to −60 mV was applied for 5.0 ms and the time to 50% decay of peak current was recorded.

###### Inactivation Time Constant (I_Na_ HEK only)

Steady-state probabilities were found at −100 mV. For voltages between −20 to 20 mV in 5 mV increments, the time to 50% decay of peak current was recorded.

##### Maximum Open Probability I_Na_ HEK and I_Na_

To constrain open probabilities, maximum open probabilities of 0.27, 0.31, 0.29 at −10, 0, 10 mV, respectively (calculated from ten Tusscher 2006(52) solved in MATLAB with ode15s) were enforced. I_to_: To best match the original I_to_ simulated current, maximum open probabilities of 0.3 and 0.45 at 25 and 50 mV, respectively were enforced (calculated from (50), solved in MATLAB with ode15s)

#### Validation set of voltage-clamp protocols

Twenty percent of experimental data was randomly chosen to serve as the validation set as is common when avoiding overfitting (63). From each curve, 80% of the data points were randomly selected and used to optimize the rate parameters. The remaining 20% of points were used to evaluate how well the model recapitulated the general trend in experimental data.

#### Action Potential Model Validation

Optimized models replaced respective currents in Tomek et al. (50) and ten Tusscher et al. (52) human ventricular action models. To simulate arrhythmogenic repetitive excitation for I_to,f_ models, an action potential was elicited by an S1 stimulus followed by an S2 stimulus at various DI intervals following S1.

### Computing Resources

All simulation code was written in C++ and containerized using Docker to run on the Amazon Web Services Batch compute cluster. Model parsing code in Python and all code is available on GitHub [https://github.com/mangoldk/AdvIonChannelMMOptimizer]. A sample K^+^ conductance voltage optimization program is also available.

## Author Contributions

KEM and JRS designed the analysis and wrote the paper. KEM executed the simulations and collected data. WW, EKJ and JMN collected, analyzed, and interpreted the human/mouse myocyte experimental data. JDM performed model validation and DB contributed to data analysis and tested the published equations.

## Classification

Biological Sciences: Biophysics and Computational Biology

## Acknowledgements

This work was supported by NIH grants R01HL136553 (JRS), R01HL143333 (JMN), R01HL150637 (JRS and JMN) and T32-HL134635 (KM). This work was also supported by an Amazon Web Services Grant (JM). The authors would also like to acknowledge Michael Southworth for his programming guidance.

